# Mass Spectrometric detection of SARS-CoV-2 virus in scrapings of the epithelium of the nasopharynx of infected patients via Nucleocapsid N protein

**DOI:** 10.1101/2020.05.24.113043

**Authors:** EN Nikolaev, MI Indeykina, AG Brzhozovskiy, AE Bugrova, AS Kononikhin, NL Starodubtseva, EV Petrotchenko, G Kovalev, CH Borchers, GT Sukhikh

## Abstract

Detection of viral RNA by PCR is currently the main diagnostic tool for COVID-19 [1]. The PCR-based test, however, shows limited sensitivity, especially at early and late stages of the disease development [2,3], and is relatively time consuming. Fast and reliable complementary methods for detecting the viral infection would be of help in the current pandemia conditions. Mass-spectrometry is one of such possibilities. We have developed a mass-spectrometry based method for the detection of the SARS CoV-2 virus in nasopharynx epithelial swabs, based on the detection of the viral nucleocapsid N protein. The N protein of the SARS-COV-2 virus, the most abundant protein in the virion, is the best candidate for mass-spectrometric detection of the infection, and MS-based detection of several peptides from the SARS-COoV-2 nucleoprotein has been reported earlier by the Sinz group [4]. Our approach shows confident identification of the N protein in patient samples even with the lowest viral loads and a much simpler preparation procedure. Our main protocol consists of virus inactivation by heating and adding of isopropanol, and tryptic digestion of the proteins sedimented from the swabs followed by MS analysis. A set of unique peptides, produced as a result of proteolysis of the nucleocapsid phosphoprotein of SARS-CoV-2, is detected. The obtained results can further be used to create fast parallel mass-spectrometric approaches for the detection of the virus in the nasopharyngeal mucosa, saliva, sputum and other physiological fluids.

## INTRODUCTION

Severe acute respiratory syndrome coronavirus 2 (SARS-CoV-2) is the causative agent of coronavirus disease 2019 (COVID-19). The virus unit consists of a single-stranded RNA with nucleoproteins enclosed within a capsid containing matrix proteins. The genome of the SARS-CoV-2 virus has been sequenced [5], and the polymerase chain reaction (PCR) method is currently the main diagnostics approach for COVID-19 diagnostics. However, it has shown limited accuracy and sensitivity, giving a high frequency of false positive and false negative results [1–3], and is relatively time consuming. The detection of viral proteins in body fluids by mass-spectrometry based methods could serve as a complementary diagnostic tool. In addition, alternative testing assays would allow us to better understand the biological activity of the virus and to suggest potential drug targets for patient treatment.

SARS-CoV-2 proteins can be grouped into two major classes - structural and non-structural proteins. Non-structural proteins are encoded by the virus, but are present only in the infected host cells and include the various enzymes and transcription factors necessary for virus replication. These proteins are not incorporated into the virion and are not as highly expressed, thus are less likely to be detected. The four structural proteins incorporated in the virion particle of SARS-CoV-2 virus are known as the S (spike), E (envelope), M (membrane), and N (nucleocapsid) proteins. The N protein encapsulates and protects the RNA genome, and the S, E, and M proteins together create the viral envelope. From the data previously obtained for SARS-CoV [6], which has a structure similar to SARS-CoV-2 structure, it can be concluded that the number of copies of E, M, and N proteins is much larger than that of S, but the E and M are relatively short and tightly membrane-bound proteins, what makes them difficult to extract, detect, and identify. Thus, the main target for mass-spectrometry based detection of SARS-CoV-2 is the N protein. The possibility of developing such methods using gargle solution samples of COVID-19 patients have been reported [4]. We have performed a pilot study on nasopharynx epithelial swabs already collected from patients with CODIV-19 for RT-qPCR and showed confident identification of the N protein of the SARS CoV-2 virus by mass-spectrometry with the use of a very basic sample preparation procedure.

Also results on the unique easily detectable peptides characteristic of the infection can further be used as targets for creating highly specific and sensitive PRM based detection methods or fast parallel mass-spectrometric approaches based on immunoprecipitation and MALDI analysis (iMALDI), allowing very fast testing for the presence of the virus in nasopharyngeal mucosa, saliva, sputum and other physiological fluids.

## MATERIALS AND METHODS

### Sample collection

All procedures for collection, transport, and preparation of the samples were carried out according to the restrictions and protocols of SR 1.3.3118-13 «Safety procedures for work with microorganisms of the I - II groups of pathogenicity (hazard)». Swabs of the mucosa of the lower part of the nasopharynx and posterior wall of the oropharynx were used for the study. The sample was collected via a sterile velor swab with a plastic applicator. The swab was introduced along the outer wall of the nose to a depth of 2-3 cm to the lower shell, and after performing a rotational movement, was removed along the outer wall of the nose. After obtaining the material, the swab (up to the place of breakage) was placed into a sterile disposable tube with transport medium, and the end of the probe was broken off. Oropharyngeal swab samples were taken with a dry sterile viscose swab by rotational movements along the surface of the tonsils, palatine arches and posterior wall of the oropharynx. In order to increase the viral concentration both the nasopharyngeal and oropharyngeal swabs were placed into a single tube, which was then sealed and marked.

The samples were collected from 5 patients with COVID-19 infection confirmed by RT-qPCR. The negative control samples were collected from 3 healthy individuals.

### Protocol for virus-inactivation of patient samples

Before the virus has been inactivated and the outside of the tubes has been disinfected, all work must be carried out in accordance with the rules of biological safety level 3 [7].

1. Transfer 0.25 mL of the specimen to a clean polypropylene 1.5-2.0 mL Eppendorf tube.
2. Inactivate by heating at 65°C for 30 minutes using a water bath or a thermostated block heater) [8]*.
3. After incubation, add 0.75 mL of 100% isopropanol to obtain a 75% solution and allow the sample to sit for 10 minutes at room temperature [9].
4. Treat the outer surfaces of the tubes with 70% isopropanol or 1% sodium hypochlorite: spill from a jet wash and let stand for 15 minutes without getting wet.

After this procedure, the virus can be considered deactivated, and the surface of the tube safe to handle.

Note: * - It is important to note, that during heat treatment, all parts of the tube are heated, so that all of the virus, even on “inaccessible” surfaces, will be inactivated.

### Sample preparation

#### Protocol 1: «standard proteomics preparation procedure»

Inactivated samples were lyophilized and resuspended in 50mM ammonium bicarbonate buffer, containing 0.1% Rapigest SF Surfactant (Waters). Reduction was carried out by incubating for 30 minutes at 50°C in 20mM DTT, followed by alkylation with 50mM iodacetamide for 45 minutes at room temperature in the dark, and tryptic digestion for 4 hours at 37°C. The reaction was terminated by adding formic acid to a final concentration of 0.5%.

#### Protocol 2: «Express preparation procedure»

Inactivated samples were cooled down to −20°C for 2 hours and centrifuged at 20 000 x g for 20 minutes. The pellet was resuspended in 50mM ammonium bicarbonate buffer, containing 0.1% Rapigest SF Surfactant (Waters) and subjected to tryptic digestion for 4 hours at 37°C. The reaction was terminated by adding formic acid to a final concentration of 0.5%.

### LC-MS/MS method

The tryptic peptides were analyzed in duplicate on a nano-HPLC Dionex Ultimate3000 system (Thermo Fisher Scientific, USA) coupled to a TIMS TOF Pro (Bruker Daltonics, USA) mass-spectrometer. The amount of sample loaded was 200 ng per injection. HPLC separation (injection volume 2 μL) was carried out using a packed emitter column (C18, 25cm x 75μm 1.6μm) (Ion Optics, Parkville, Australia) [10] by gradient elution. Mobile phase A was 0.1% formic acid in water; mobile phase B was 0.1% formic acid in acetonitrile. LC separation was achieved at a flow of 400 nL/min using a 40-min gradient from 4% to 90% of phase B.

Mass spectrometric measurements were carried out using the Parallel Accumulation Serial Fragmentation (PASEF™) [11] acquisition method. The ESI source settings were the following: 4500 V capillary voltage, 500 V end plate offset, 3.0 L/min of dry gas at temperature of 180oC. The measurements were carried out over the m/z range from 100 to 1700 Th. The range of ion mobilities included values from 0.60-1.60 Vs/cm2 (1/k0). The total cycle time was set at 1.16 sec and the number of PASEF MS/MS scans was set to 10. For low sample amounts, the total cycle time was set to 1.88 sec.

### Data Analysis

The obtained data was analyzed using PEAKS Studio 8.5 and MaxQuant version 1.6.7.0 using the following parameters: parent mass error tolerance - 20ppm; fragment mass error tolerance - 0.03Da. Due to light denaturation conditions, the absence of reduction and alkylation steps in one of the sample preparation approaches and short hydrolysis time - up to 3 missed cleavages were allowed, but only peptides with both trypsin-specific ends were considered. Oxidation of methionine and carbamidomethylation of cysteine residues were set as possible variable modifications and up to 3 variable modifications per peptide were allowed. The search was carried out using the Swissprot SARS-COV-2 database with the human one set as the contamination database. FDR thresholds for all stages were set to 0.01 (1%) or lower.

## RESULTS

Approximately 1000-1500 proteins were identified in each sample, among which the P0DTC9|NCAP_SARS2 Nucleoprotein of Severe acute respiratory syndrome coronavirus 2 was registered (Figure 1).

**Figure 1.**
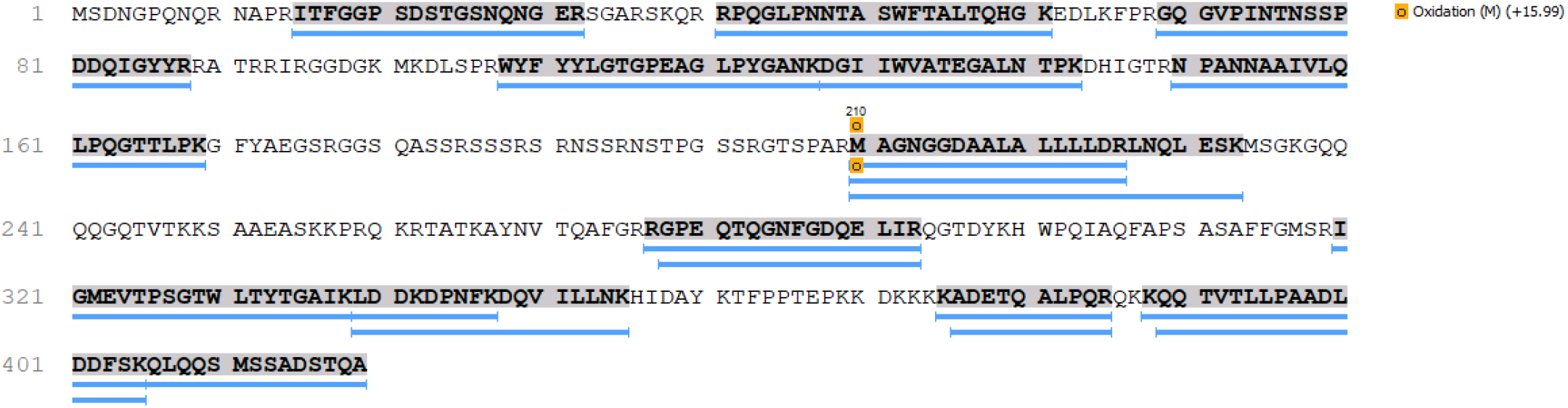
Sequence coverage of the P0DTC9|NCAP_SARS2 Nucleoprotein from SARS CoV-2.

**Figure 2.**
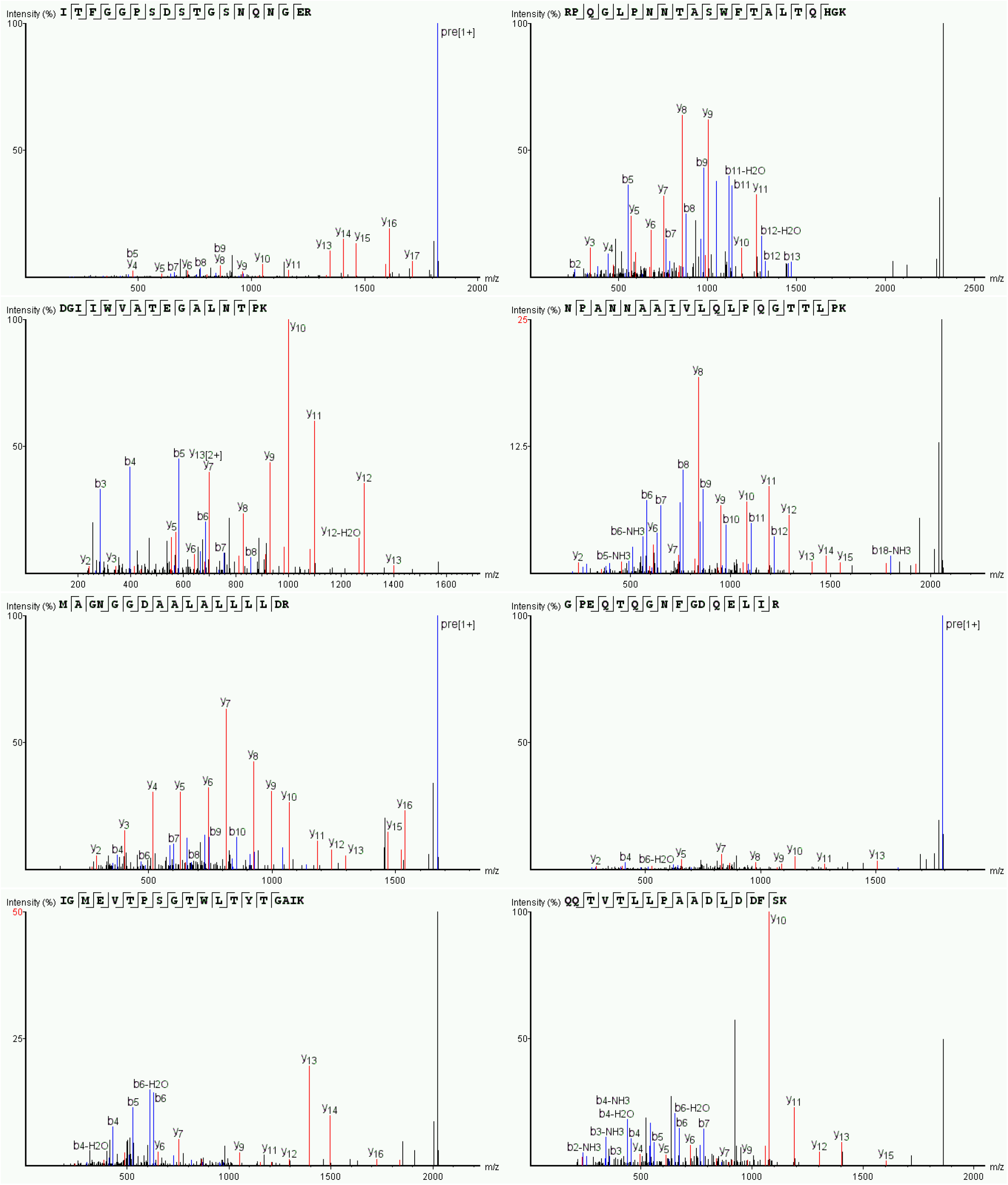
Examples of MS/MS data of detected peptides of **P0DTC9|NCAP_SARS2 Nucleoprotein**.

Depending on the viral content in the samples, preparation protocol and processing software used 1-17 peptides of the N protein were reliably detected and identified in the COVID-19 patient samples (Tables 1 and 2).

**Table 1.**
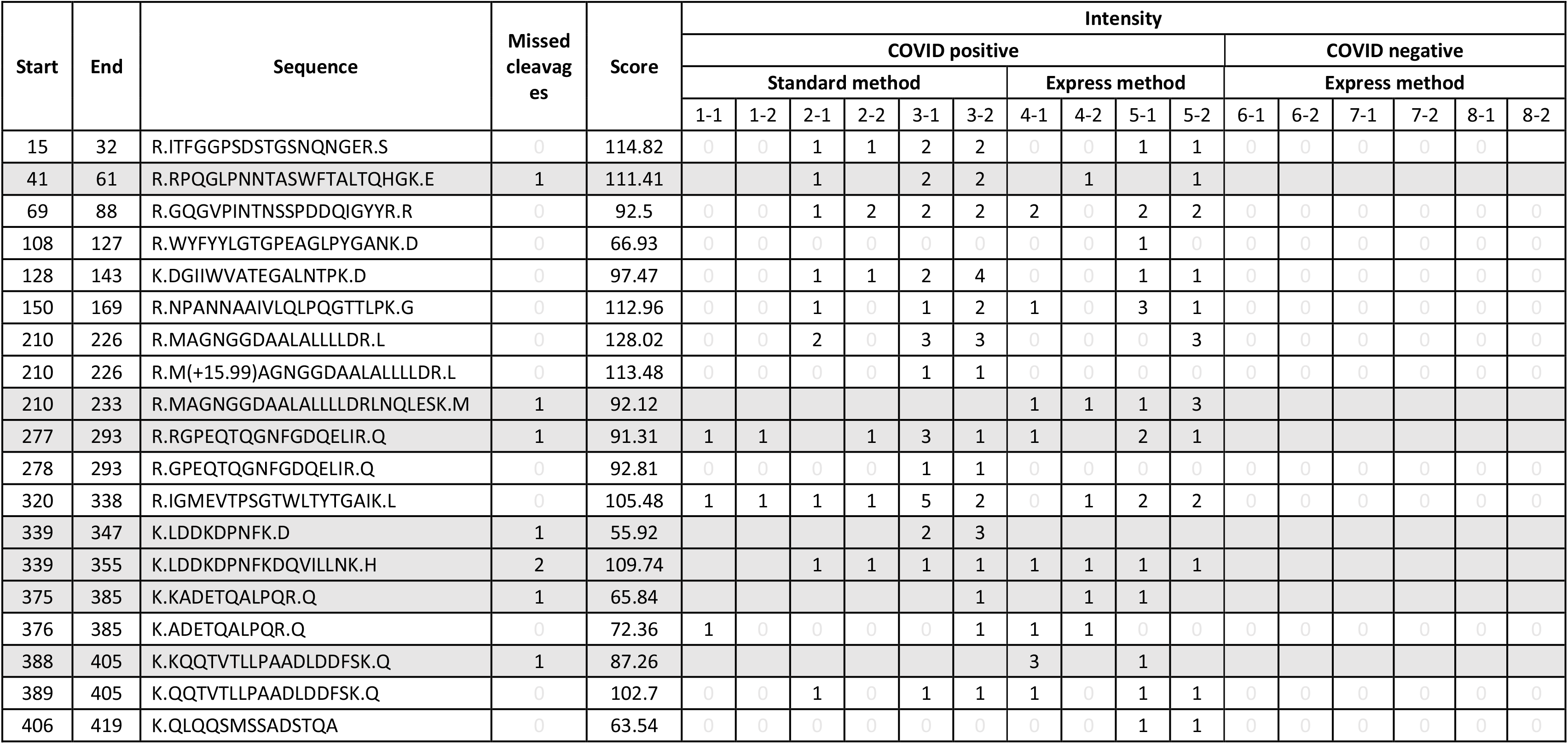
Peptides from the P0DTC9|NCAP_SARS2 Nucleoprotein identified via PEAKS Studio in different samples sample preparation protocol. Samples 1-5 were collected from 5 patients with COVID-19 (1-3 were prepared by Protocol 1; 4-5 by Protocol 2). Samples 6-8 – negative control (healthy individuals). Two LC MS/MS runs were performed for each sample. The numbers in the table correspond to spectral counts for each peptide.

**Table 2.**
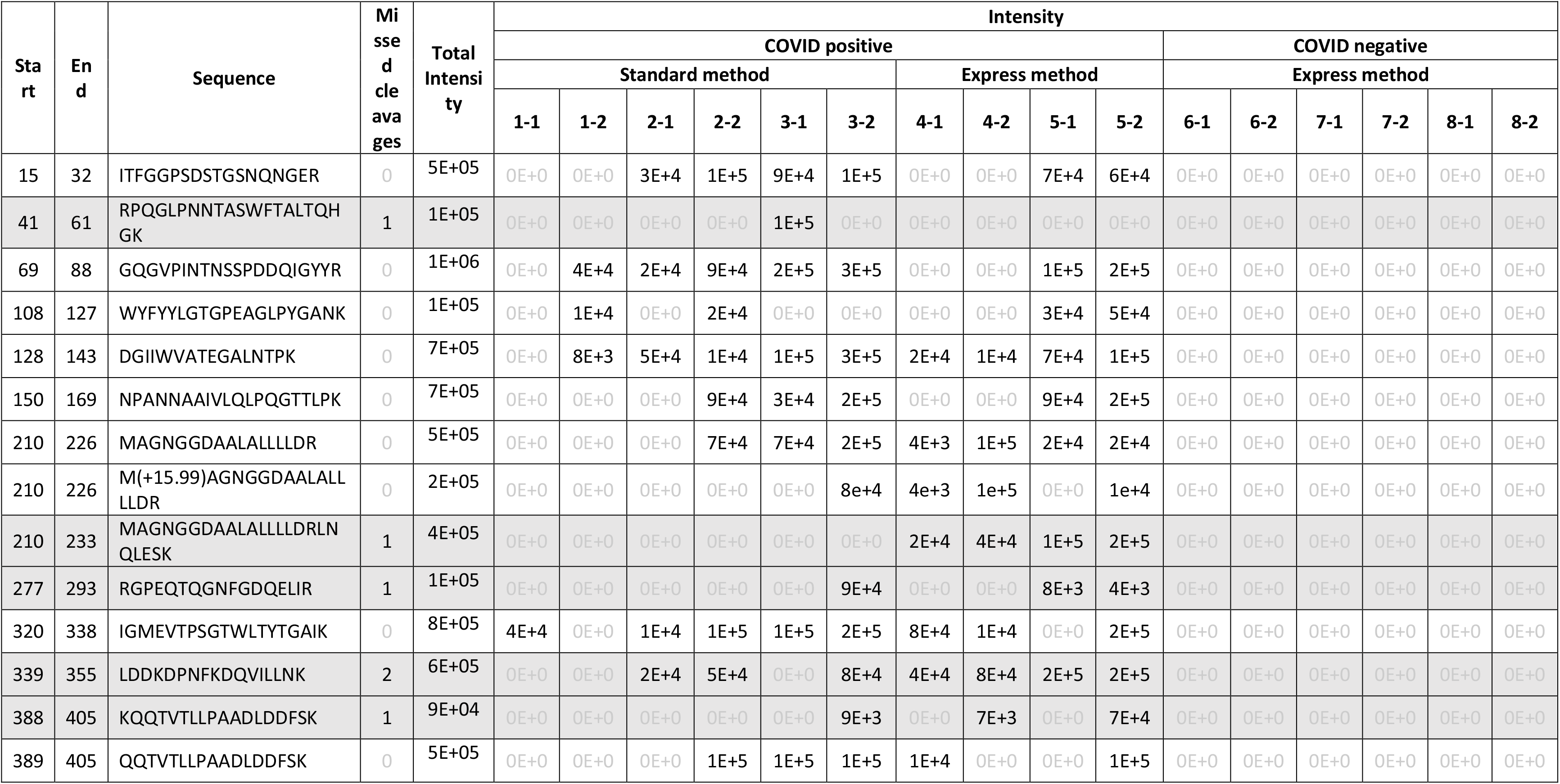
Peptides from the **P0DTC9|NCAP_SARS2 Nucleoprotein** identified **via MaxQuant** in different samples depending on sample preparation protocol. **Samples 1-5** collected from 5 patients with COVID-19 (1-3 were prepared by Protocol 1; 4-5 by Protocol 2**). Samples 6-8** – negative control (healthy individuals). Two LC MS/MS runs were performed for each sample. The numbers in the table correspond to the intensities of each peptide.

The N protein, being the most abundant protein in the virion, is the best candidate for mass-spectrometric detection of the infection, and so its detection is expectable. Also MS-based detection of several peptides from the SARS-COoV-2 nucleoprotein has been reported earlier by the Sinz group [4] in the gargle solution samples of COVID-19 patients. We have performed a pilot study on nasopharynx epithelial swabs already collected from patients with CODIV-19 for RT-qPCR and showed confident identification of the N protein with the use of a very basic sample preparation procedure.

More than that the express procedure allowed better detection of the N protein than the more thorough one, standardly used for proteomic analysis. This is probably due to the significantly lower amounts of peptides from the much more abundant host proteins, which require reduction, alkylation, deglycosylation and other preparative steps for good sequence coverage.

Also since an untargeted LC-MS/MS with data-dependent acquisition approach was used to identify as many proteins as possible, it is expected that the use of targeted approaches aimed at monitoring the presence of this exact protein will result in significantly lower detection limits. For example the use of PRM approaches on triple-quadrupole mass-spectrometers basing on the peptides identified in this study may allow to go below the sensitivity of RT-qPCR, while application of immunoprecipitation methods with subsequent MALDI analysis or even MALDI detection of viral proteins directly on the immobilized antibodies (iMALDI) will allow to shorten the processing times to as low as 1hour.

The observation of over 1000 host proteins also suggests that this approach could be used for detecting and studying the changes caused by the viral infection in the proteome of host cells, as well as the response of the organism to these conditions.

## Notes

### Competing Interest Statement

The authors have declared no competing interest.

